# GeneArbiter improves plant genome annotations while preserving established gene models

**DOI:** 10.64898/2026.11.26.769510

**Authors:** Xiaoxiao Wang, Zhanxiang Zong, Shaochen Jia, Rui Zhong, Hu Zhao

**Affiliations:** National Key Laboratory of Crop Genetic Improvement, Hubei Hongshan Laboratory, Huazhong Agricultural University, Wuhan 430070, China

## Abstract

As plant genome resources expand, updating established annotations requires incorporating improved gene models without disrupting correct structures or stable identifiers. We developed GeneArbiter, an evidence-guided framework that treats annotation updating as a correction–preservation problem. GeneArbiter uses the current annotation as a backbone, integrates alternative models with RNA splice junction and homologous-protein evidence, and applies constrained language-model arbitration followed by deterministic release and identifier reconciliation. In rice MH63, GeneArbiter recovered 4,701 reference loci missed by the current annotation while losing 1,890 current-exact loci, yielding a net gain of 2,811 exact matches and increasing the gene-level exact-hit rate from 70.77% to 77.77%. It retained 97.54% of protein-coding genes in the mature TAIR12 annotation and improved protein-set quality metrics across 16 plant genomes. Independent transcript and protein evidence further supported GeneArbiter-selected structural revisions at reference-discordant loci. These results establish GeneArbiter as a conservative and traceable framework for maintaining plant genome annotations as new predictions and biological evidence accumulate.

## Results

As PLANeT and other plant genome initiatives expand high-quality reference genomes across plant lineages, maintaining accurate and stable genome annotations is increasingly important for gene discovery, comparative genomics and functional studies (Wang et al., 2026). Improved *de novo* prediction yields better candidates, while growing transcriptomic and homologous-protein data provide orthogonal evidence for evaluating alternative gene structures (Holst et al., 2026; Zhang et al., 2026; Gabriel et al., 2024). However, generating a new consensus annotation is not equivalent to updating an established release. Evidence-integration methods such as EVidenceModeler (EVM) are primarily designed to construct consensus gene sets, whereas updating released versions must incorporate improved models without unnecessarily replacing correct existing structures, disrupting stable identifiers or indiscriminately expanding gene catalogues (Haas et al., 2008). Recent updates illustrate this distinction: ZS11 transferred legacy models and added *de novo* predictions to an improved assembly, whereas TAIR12 preserved compatibility with prior annotations and identifiers on a new assembly through community expert review (Xu et al., 2026; Reiser et al., 2026).

We therefore developed GeneArbiter, an automated framework for evidence-guided annotation updating that uses the current annotation as a backbone and selectively incorporates better-supported candidates (Figure 1A). In MH63, the current annotation exactly matched 28,405 of 40,140 curated protein-coding genes at the complete CDS-chain level, yet five *de novo* annotations failed to recover 10,600 of these, and all six annotations lacked an exact candidate for 4,692 curated genes (Figure 1B). These results illustrate why improved *de novo* prediction alone does not solve the annotation-update problem: new predictions can recover missed structures, but wholesale replacement can simultaneously discard many correct existing models. GeneArbiter addresses this correction– preservation challenge by grouping overlapping and structurally connected models into dynamic loci, integrating RNA splice-junction and protein-homology evidence into structured evidence cards, and performing deterministic structural checks before arbitration. Candidate sources are anonymized; a constrained language model is restricted to selecting among predefined candidate sets rather than generating gene structures or genomic coordinates, after which deterministic software applies release policies, reconciles identifiers and exports GFF3 annotations with traceable mappings and provenance records.

**Figure 1.**
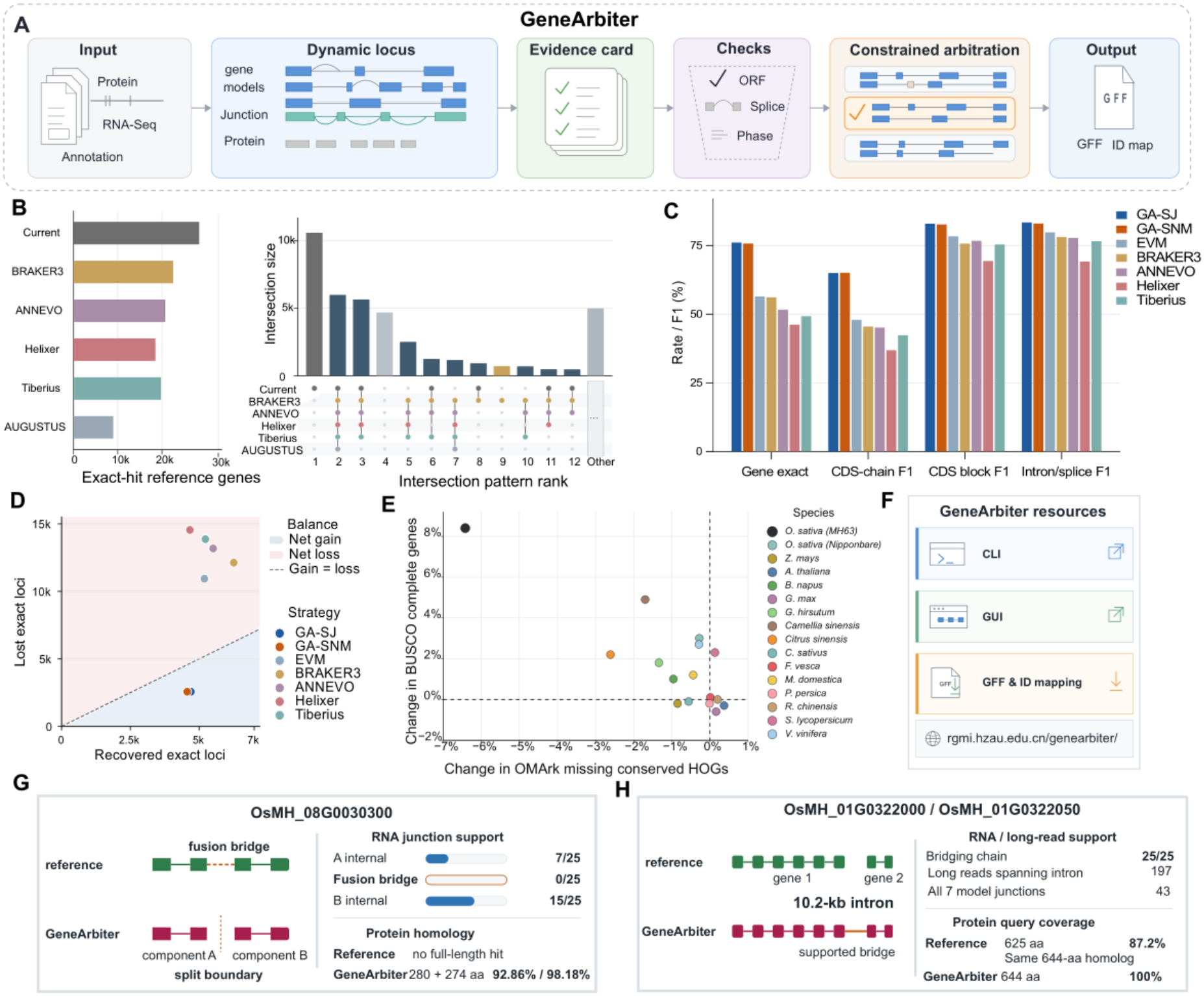
GeneArbiter updates established plant genome annotations while balancing correction and preservation. **(A)** Workflow. Existing and candidate annotations are organized into dynamic loci together with RNA splice-junction and protein-homology evidence. Structured evidence cards and deterministic ORF, splice and CDS-phase checks inform constrained arbitration among listed candidate sets; deterministic software exports GFF3 annotations and identifier mappings. **(B)** Exact CDS-chain overlap among the current MH63 annotation and five *de novo* annotations (BRAKER3, ANNEVO, Helixer, Tiberius and AUGUSTUS), evaluated against 40,140 manually curated protein-coding genes. Horizontal bars show reference genes with at least one exact transcript. Intersection bars and the matrix show the 12 largest membership patterns; the remaining 52 patterns are pooled as Other (4,994 genes). An exact hit requires the complete CDS interval chain, chromosome and strand to match a reference transcript. **(C)** Whole-genome MH63 benchmark against the same curated reference, comparing GeneArbiter-SJ (GA-SJ) and GeneArbiter-SNM (GA-SNM) with EVM, BRAKER3, ANNEVO, Helixer and Tiberius. Gene exact-hit rate uses 40,140 reference protein-coding genes as its denominator; the other metrics are exact CDS-chain, CDS-block and intron/splice F1 scores. **(D)** Gain–loss balance relative to the current annotation. Recovered exact loci are reference genes missed by the current annotation but exactly recovered by each strategy; lost exact loci are genes exactly matched by the current annotation but no longer exactly matched by that strategy. The dashed line indicates equal gains and losses, with points below the line representing a net gain. **(E)** Paired changes in BUSCO complete percentage and OMArk missing conserved-HOG percentage across 16 plant genomes, calculated as GA-SJ minus the original annotation in percentage points. Positive BUSCO changes and negative OMArk missing-HOG changes indicate improvement in these quality proxies, which do not independently measure gene-structure accuracy. **(F)** Software and data availability. The GeneArbiter v0.4.0 software release is available on GitHub; the project download site provides GA-SJ and GA-SNM annotations and identifier mappings for all 16 genomes. **(G)** Split case at OsMH_08G0030300. The reference contains a fused model, whereas GeneArbiter selects two component models. RNA-seq junction support is present within component A (7/25) and component B (15/25) but absent across the fusion bridge (0/25). No single full-length homolog supports the reference fusion, whereas the two GeneArbiter proteins (280 and 274 amino acids) match separate homologs with 92.86% and 98.18% identity, respectively. **(H)** Merge case at OsMH_01G0322000/OsMH_01G0322050. All seven junctions of the merged model, including the cross-gene junction, were detected in all 25 short-read RNA-seq runs; 197 long reads spanned the 10.2-kb intron, including 43 containing all seven model junctions. BLASTP against the same 644-aa Nipponbare protein (Os01t0527400-01) gave 87.2% query coverage for the 625-aa reference protein and 100% for the 644-aa merged protein.

Using the GSAman-refined MH63 annotation, which was withheld from arbitration and used only as a held-out evaluation reference (Chen et al., 2026), we evaluated GeneArbiter genome-wide. For loci absent from the current annotation, the single-tool junction-supported (SJ) policy accepts a selected coding structure predicted by at least two tools, or by one tool with splice-junction support; the more conservative supported-novel multi-tool-only (SNM) policy excludes the latter route. In MH63, SJ recovered 40.06% (4,701) of reference loci not exactly matched by the current annotation while retaining 93.35% of current-exact loci, yielding 4,701 recovered versus 1,890 lost exact hits and a net gain of 2,811 (Figure 1D). EVM recovered 5,191 current-missed loci but lost 10,931 current-exact loci, a net loss of 5,740; BRAKER3, ANNEVO, Helixer and Tiberius similarly produced net losses of 5,867–9,883. GeneArbiter therefore recovered missed structures while limiting disruption of correct models. This balance raised the gene-level exact-hit rate from 70.77% to 77.77%, the highest among seven methods; the five external methods reached 46.14–56.46% (Figure 1C). Its exact CDS-chain F1 was 65.47%, versus 36.90–47.93% for external methods, with consistent improvements at CDS-block and intron levels. To test preservation in a mature annotation, we used community-curated, expert-reviewed TAIR12 as a fixed reference (Reiser et al., 2026), conservatively treating any original structure not exactly retained as a potential unintended change. GeneArbiter exactly retained 97.54% of protein-coding genes and 91.86% of coding transcripts, with an exact CDS-chain F1 of 94.32% and a 2.46% gene-level upper bound for potential unintended structural changes (Supplemental Figure S1). SNM similarly reached a 77.43% gene-level exact-hit rate in MH63 while retaining the same TAIR12 gene and transcript proportions.

We next distinguished limits imposed by candidate availability from arbitration’s ability to identify the correct structure within that space. Among 19,009 MH63 cards overlapping the reference, 16,941 (89.12%) contained a reference-exact candidate, defining the recoverable ceiling under the current design. GeneArbiter selected a set containing such a model in 14,662 cards (77.13% overall), an 86.55% success rate when an exact solution was available. Among 9,874 cards containing both exact and non-exact candidates, reference-exact selection reached 76.60%, versus a candidate-composition-adjusted random expectation of 50.01% (Supplemental Figure S2). We also compared constrained language-model arbitration with seven deterministic selectors on the same candidate sets. Deterministic rules made unique calls for only 3.02–82.77% of API-routed loci, whereas the language model returned a valid listed choice at every locus. Where a rule made a unique call, disagreed with the language model and a reference-exact candidate existed, the language model more often selected the reference-exact set for all seven rules, reaching 84.02% versus 61.19% for the score-trace rule (Supplemental Figure S3).

Ablation analyses showed that both candidate diversity and biological evidence contributed to performance. Because dynamic loci were rebuilt for each candidate-pool condition, the number of evaluated current-error loci changed from 251 with ANNEVO alone to 267 after adding Helixer and remained 267 after adding Tiberius; this was therefore a full-workflow, condition-specific comparison rather than a paired comparison over a fixed locus set. Across these conditions, loci containing a reference-exact candidate increased from 113/251 (45.02%) to 133/267 (49.81%), and loci whose final selection recovered one increased from 91/251 (36.25%) to 106/267 (39.70%). With the three-tool candidate pool fixed, removing RNA or protein evidence reduced recovery efficiency among loci with a reference-exact candidate from 79.70% to 66.17% or 77.44%, respectively, indicating a larger RNA contribution in this MH63 subset (Supplemental Figure S2).

To assess gene-level run-to-run robustness, we applied the same SJ release workflow to three frozen MH63 short-read-only whole-genome runs, which yielded gene-level exact-hit rates of 77.77–78.09%. Two independent runs under matched V4.1 settings differed by only 0.08 percentage points and agreed on 97.85% of candidate-set selections (Supplemental Figure S4). Together, these analyses show that GeneArbiter’s performance depends on the availability of complementary candidates and effective integration of orthogonal biological evidence while remaining reproducible across runs.

To assess portability beyond MH63, we applied GeneArbiter to 16 plant genomes without species-specific language-model training. Because most lacked independent structure-level references, paired changes in BUSCO completeness and OMArk missing HOGs were treated as protein-set quality proxies, not direct measures of gene-structure accuracy. Original annotations already averaged 95.6% BUSCO completeness, with 12 genomes at 95% or higher. Against these baselines, SJ increased BUSCO completeness by an unweighted mean of 1.64 percentage points and reduced the proportion of missing conserved HOGs measured by OMArk by 0.91 percentage points (Figure 1E). BUSCO improved in 10 genomes and OMArk in 11. Excluding MH63, whose BUSCO increased from 88.7% to 97.1%, mean BUSCO improvement across the other 15 genomes remained 1.19 points. SNM showed similar trends (Supplemental Figure S5).

At two MH63 loci counted as errors by exact-match evaluation against the manually refined reference, independent RNA and protein evidence instead supported GeneArbiter’s structural choices. At OsMH_08G0030300, the reference fused two components, but their internal junctions were detected in 7/25 and 15/25 RNA-seq runs, whereas the fusion bridge was not detected in any run (0/25). The two GeneArbiter proteins matched distinct near-full-length homologues with 92.86% and 98.18% identity, supporting the split (Figure 1G). At OsMH_01G0322000/OsMH_01G0322050, GeneArbiter selected a single model spanning a repeat-rich 10.2-kb intron rather than two reference genes. All seven junctions of the merged model, including the cross-gene junction, were detected in all 25 short-read RNA-seq runs; 197 long reads spanned the 10.2-kb intron, including 43 containing all seven model junctions. Against the same Nipponbare homologue, query coverage was 87.2% for the 625-aa reference protein and 100% for the 644-aa merged protein (Figure 1H). Detailed IGV and run-level junction-support views for both cases are provided in Supplemental Figure S6. The split recovered two tandem DUF26 cysteine-rich secretory-protein genes conserved across *Oryza*, whereas the merge reconstructed a full-length BAH–TFIIS-family protein across the long intron. Additional gene-addition, splitting and merging examples are shown in Supplemental Figure S7. These orthogonal validations indicate that some choices counted as errors in reference-dependent evaluation may instead be biologically supported structural corrections and that individual models in manually refined annotations can still warrant re-examination.

API usage and direct costs for the 16-genome analysis are reported in Supplemental Figure S8 and Source Data; costs exclude upstream computation. The GeneArbiter v0.4.0 software release is available on GitHub (https://github.com/wxx7556923/GeneArbiter/releases/tag/v0.4.0), while SJ and SNM annotations and identifier mappings for all 16 genomes are available at https://rgmi.hzau.edu.cn/genearbiter/ (Figure 1F). The local graphical interface and web download portal are shown in Supplemental Figure S9A and S9B, respectively. Together, our results show that GeneArbiter enables conservative, traceable updates to established plant genome annotations. Rather than replacing annotations wholesale, it makes model-level decisions explicit and evidence-based, incorporating better-supported structures while preserving reliable existing models and continuity across versions.

## Limitations

GeneArbiter is designed to refine established genome annotations rather than to replace de novo gene prediction. Its performance is therefore limited by the availability and quality of candidate gene models and supporting biological evidence. Although repeated runs showed stable genome-wide performance, language-model arbitration can introduce locus-level variability and may be affected by changes in model versions. Furthermore, reference-based exact-match evaluation assumes that the reference structures are correct, whereas BUSCO and OMArk measure complementary aspects of protein-set quality rather than individual gene-structure accuracy. Further evaluation using independent curated annotations and alternative arbitration strategies will help define the applicability and limitations of the framework.

## Materials and methods

Detailed descriptions of genome annotation resources, candidate model construction, dynamic locus definition, biological evidence integration, constrained language-model arbitration, release policies, identifier reconciliation, and benchmarking procedures are provided in the Supplemental Methods.

## Data and code availability

The public RNA-seq accessions and the assemblies and baseline annotations used in this study are listed in Supplemental Tables S1 and S2. Numerical data and locus-level information underlying main and supplemental figures, together with recorded API token usage for the 16-genome analyses, are provided as Source Data. Genome assemblies, baseline annotations, GeneArbiter-SJ and GeneArbiter-SNM GFF3 outputs, and identifier mapping files for all 16 genomes are publicly available at https://rgmi.hzau.edu.cn/genearbiter/.

GeneArbiter v0.4.0 source code is available from the GitHub release (https://github.com/wxx7556923/GeneArbiter/releases/tag/v0.4.0), and downloadable command-line and Windows GUI packages are provided in the tagged downloads directory (https://github.com/wxx7556923/GeneArbiter/tree/v0.4.0/downloads). The version used in this study is permanently archived on Zenodo (https://doi.org/10.5281/zenodo.23113893). GeneArbiter is released under the MIT License.

## Acknowledgments

We acknowledge the support of the high-performance computing platform at the National Key Laboratory of Crop Genetic Improvement at Huazhong Agricultural University.

## Author contributions

H.Z. designed the study, supervised the research, and drafted and revised the manuscript. X.X.W. conducted the analyses, developed the toolkit, and wrote and revised the manuscript. Z.X.Z., S.C.J. and R.Z. conducted the analyses and developed the toolkit. All authors read and approved the final manuscript.

