## Supplementary Information for "GeneArbiter improves plant genome annotations while preserving established gene models"

### Methods

#### Reference annotations, candidate models and evidence resources

GeneArbiter takes a genome assembly, its existing protein-coding annotation (the current backbone), one to three candidate gene annotations and optional RNA and homologous-protein evidence. We analysed 16 plant genomes representing 15 species with their corresponding assembly and baseline annotation versions (Supplemental Table S1). The principal structural benchmark used the MH63RS3 rice assembly, its MHRS3_nonTE annotation and candidate models from ANNEVO, Helixer and Tiberius. These three tools were run with the pretrained plant or angiosperm models listed in Supplemental Table S3; their GFF files were normalized to a common gene–transcript–exon/CDS hierarchy before arbitration, whereas the current annotation was retained as the backbone. BRAKER3, AUGUSTUS and EVM served as comparators rather than additional sources in the principal three-tool GeneArbiter run. The curated MH63 annotation was not supplied to candidate generation, evidence cards or the language-model API. Figure 1B compares the current annotation with BRAKER3, ANNEVO, Helixer, Tiberius and AUGUSTUS.

#### RNA-seq processing and splice evidence

RNA-seq samples, tissue labels, library types and run or sample identifiers are listed individually in Supplemental Table S2. The principal MH63 run used 25 Illumina paired-end runs as short-read evidence. Five MH63 long-read runs (two ONT cDNA, two ONT direct RNA and one PacBio Iso-Seq) were used for the independent locus-level checks in Figure 1G and 1H; they were not supplied to the three whole-genome runs in Supplemental Figure S4. For each of the other 15 genomes, three to nine Illumina RNA-seq runs or libraries were used as splice-junction evidence; dataset identifiers, tissues and replicate metadata are provided in Supplemental Tables S1 and S2.

MH63 raw reads were quality-checked with FastQC v0.11.9. Selected short-read evidence was aligned to the matching genome with HISAT2 v2.2.1 using --dta without annotation-derived exon or splice-site hints. Alignments were sorted and indexed with SAMtools v1.23.1. Splice junctions were extracted from mapped primary alignments; samtools view -F 2308 excludes unmapped, secondary and supplementary alignments, but does not exclude reads carrying the duplicate flag. Junction coordinates and supporting read and sample counts were aggregated across selected runs. During evidence-card construction, a junction required at least three supporting reads from at least one sample. Each candidate intron was classified by exact-coordinate support, nearby alternative junctions and missing observations; absence of a detected junction alone was not treated as proof that a transcript was incorrect. Model-level coverage summaries were generated, but coverage-derived risk/protection fields were excluded from the compact cards used in the evaluated decisions.

For the independent locus-level checks in Figure 1G and 1H, MH63 long reads were aligned to the genome with minimap2 v2.27-r1193 using splice-aware presets: -ax splice -k14 --secondary=no --MD for ONT cDNA, -ax splice -uf -k14 --secondary=no --MD for ONT direct RNA, and -ax splice:hq -uf --secondary=no --MD for PacBio Iso-Seq. Primary mapped alignments were parsed for CIGAR N operations, and ordered intron chains were compared with the tested model. We counted reads spanning the cross-gene intron, reads carrying all seven junctions of the merged model and reads covering its complete coding span. These long-read observations were used for the Figure 1G/H locus checks, not as input to the three S4 whole-genome decisions.

#### Homologous-protein evidence

The MH63 main analysis used the public poaceae_public_v1 protein library: 386,868 longest-per-gene sequences from nine grass species, including public *Oryza sativa*, *O. glaberrima* and *O. brachyantha* proteins, downloaded from Ensembl Plants release 116. The library excluded proteins derived from the manually curated MH63 benchmark annotation. Source sequences were filtered for length of at least 30 amino acids, valid amino-acid symbols, at most 20% unknown (X) residues, annotated protein-coding biotypes where available and obvious transposon-like headers; one longest protein was retained per species and gene. For other target genomes, homolog-library composition and target-species exclusions are listed in Supplemental Table S1. The PN40024 analysis used public *Vitis* peptides independent of the T2T annotation evaluated here.

Protein sequences were aligned to each target genome with miniprot v0.18-r281, and the resulting protein-alignment GFF was supplied as evidence rather than as another candidate gene annotation. Cards recorded overlapping protein hits, best target and reported identity. The numerical protein-support component described below counts candidate models with an overlapping protein-alignment feature; this overlap is a support heuristic and does not establish a full-length, structurally exact orthologue.

#### Dynamic loci, evidence cards and candidate scores

GeneArbiter grouped overlapping or structurally connected current and candidate models into dynamic loci. RNA junction and protein alignments could connect nearby model components; these loci were defined by local evidence relationships rather than a single fixed genomic window. The evaluated MH63 configuration set 50-kb soft and 100-kb hard limits for long AI-facing loci and capped the number of current genes and models per source in a long window at three each. Within a locus, the pipeline enumerated competing annotation sets, collapsed structurally equivalent choices into AI-facing groups and summarized CDS presence, ORF and phase checks, RNA junctions, optional long-read chains, protein overlaps, component coverage, split/merge topology and explicit risks. Candidate-source identities were anonymized in the compact card. Numerical scores were used for candidate ranking; the final AI-facing card showed categorical evidence and risk summaries, without raw numerical scores or source identities.

For each candidate set, a deterministic evidence-support score was assembled from five nonnegative components (maximum raw subtotal 95), followed by risk penalties and clipping to 0–100. The component rules were:

| Component | Maximum | Scoring rule |
|---|---|---|
| Coding QC | 25 | 25 if at least 95% of transcripts pass; 17 if at least 60% pass; 8 if some pass; 0 if none are assessable. If any transcript fails, use max(0, 8 − 8 × fail_count/transcript_count) instead. |
| Short-read junctions | 25 | 25 if at least 95% of transcripts have full junction support; 18 if at least 80% have full or partial support; 9 if any have such support; 5 if unassessable with no none_observed call; otherwise 0. |
| Long-read chains | 15 | When configured: 15 if at least 80% of transcripts have exact-chain support; 10 if any have an exact chain or at least 80% have exact/partial support; 5 if some have partial support; 3 for unassessable single-exon models without unsupported transcripts; otherwise 0. |
| Homologous protein | 20 | 20 if at least 80% of transcripts overlap a protein hit; 10 if some do; otherwise 0. |
| Structural flags | 10 | Add 4 for CDS present, 4 for all introns junction-supported and 2 for model-level full intron-chain support, capped at 10. |

Risk penalties were subtracted for validation failure (min[25, 12 + 5 × fail_count]), warnings (min[8, 3 × warning_count]), conflicting short-read junctions (min[15, 5 × conflict_count]), a conflicting long-read chain (5), and distinct raw risk tags (min[10, 2 × tag_count]). With run-level normalization, only evidence classes configured for the entire run entered the positive subtotal: coding QC and structural flags always remained active; short reads, long reads and protein evidence were activated by their configured input types. If $W$ is the sum of their active maximum weights, the score was clip[(active positive subtotal) × 95/W + risk penalty, 0, 100]. Penalties were not scaled. The principal MH63 short-read-only run had $W = 80$ because no long-read evidence was configured. A locus with no observed support still belonged to a configured evidence class and did not trigger locus-specific renormalization. This score was not visible as a number to the language model and was not a benchmark accuracy estimate or a mandatory highest-score decision; agreement among candidate sources did not add numerical score points.

#### Constrained language-model arbitration and annotation release

Cards requiring API arbitration were sent to the deepseek-chat model alias through an OpenAI-compatible chat-completions endpoint in JSON-object mode. The evaluated MH63 configuration used temperature 0, a maximum of 8,192 output tokens, a 240-s timeout, at most two attempts and 60 workers. The original Chinese system instruction and per-card user template are reproduced verbatim in the **Prompt appendix** of the Supplementary Information. The instruction requires the model to choose a set_id already listed in the card, prohibits generation of genomic coordinates or GFF records, and limits risk_flagged to unresolved mutually exclusive sets; even a flagged response must select a provisional listed set. Returned JSON was checked for the card identifier, allowed fields and values, a listed selected_set_id and requires_validation=true. Candidate selection did not use the curated reference annotation.

To audit direct API usage, we summed the usage fields in the raw provider responses of the completed decision run for each of the 16 genomes, including responses from retry attempts. Prompt, completion, cache-hit and cache-miss tokens are reported separately in Source_Data/Sixteen_genome_API_token_usage.tsv; all 16 completed runs had usage records for every saved response. Separately, the RUN2_V4.1 MH63 technical run had 19,928 response records totaling 47,830,228 tokens, exactly matching the request and token totals shown in the Figure S8 dashboard screenshot. Its displayed CNY 45.22 charge is an observed dashboard value, not a price-derived estimate. These figures cover API inference only; evidence preparation, candidate prediction, alignment, local computation and exploratory runs are outside this accounting.

The selected models were passed through deterministic support and release rules, ID reconciliation and GFF3 quality control. The current annotation remained the backbone; a rejected replacement did not remove its current gene blocks. For loci absent from the current annotation, the SJ release policy accepted an AI-selected coding structure supported by at least two candidate tools, or a single-tool structure with splice-junction support and the required evidence/risk status. SNM omitted the single-tool route. Stable current IDs were reused where possible; novel, split and merge events received traceable IDs and gene/transcript mapping records. Release GFF3 files and compact provenance traces were exported without another language-model call or access to the manually curated MH63 reference.

##### Structural benchmarks, ablations and quality proxies

For MH63, a manually refined annotation was used as the evaluation reference. A protein-coding reference gene was an exact hit when at least one predicted transcript on the same chromosome and strand had an identical ordered chain of complete CDS intervals to one reference transcript. Gene exact-hit rate used 40,140 reference coding genes as the denominator; an exact miss did not imply absence of the entire gene. Figure 1B grouped per-gene exact-hit indicators for the six displayed annotations, ranked their intersections and combined patterns outside the largest 12 into Other. Figure 1C compared GeneArbiter-SJ and -SNM with EVM, BRAKER3, ANNEVO, Helixer and Tiberius using the same reference and evaluator.

The BRAKER3 comparator used v3.0.8 in ETP mode with RNA-seq and homologous-protein evidence on the soft-masked MH63RS3 genome. The evaluated GFF3 used GeneMark-ETP intermediate predictions in the AUGUSTUS step (--skipGeneMark-ETP). AUGUSTUS v3.3.3 began with 21,107 training genes, retained 21,056 after structural filtering and randomly sampled 8,000 before redundancy filtering, leaving 7,796 genes for training. Its parameter optimization used five rounds and eightfold cross-validation. For EVM v2.1.0, the fixed source weights were 2 for the current annotation and 1 each for ANNEVO, Helixer, Tiberius, miniprot protein alignments and unguided short-read StringTie transcript alignments. The EVM run used 500-kb segments, 10-kb overlaps and 16 CPUs; these weights were supplied as input parameters rather than learned by EVM (Supplemental Table S3). Exact CDS-chain, CDS-block and intron/splice F1 values were calculated as 2 × exact matches/(prediction total + reference total) for each feature unit.

To evaluate the decision layer (Supplemental Figure S3), seven deterministic selectors operated on the same fixed AI-facing candidate groups at 19,928 API-routed MH63 loci. A rule had a unique call only when one group achieved its best score; ties were no-calls. Decision coverage was unique calls divided by 19,928. For each rule, selector success was assessed only where the rule and AI disagreed and at least one reference-exact candidate set was available; the numerator was the number of selected sets containing at least one reference-exact model, and each rule had its own denominator. The rule definitions were: SJ-only scores of 4, 3, 2, 1 and 0 for full, partial, limited, conflicting and absent/unassessable junction evidence; protein-only scores of 1, 0.5 and 0 for supported, partial and absent/unconfigured evidence; minimum risk −(risk-tag count + 3 × low-priority-extra indicator + 2 × unsupported-component indicator); and component fit 3 × core/complete − 2 × unsupported component + biologically supported − topology risk + 0.2 × supported-component count − 0.8 × unsupported-extra count − 0.8 × missing-strong count. The two composite rules used 3 × SJ + component + risk, with the protein-aware rule additionally adding 2 × protein. Score-trace top used the internal candidate rank score, or the normalized score if absent; after structural-equivalence collapse, a group received its maximum member score. These rules used no curated-reference labels during selection. Additional analyses compared selection with a candidate-composition-adjusted random expectation, incrementally added ANNEVO, Helixer and Tiberius in a separate 500-gene sample, removed RNA or protein evidence with the three-tool pool fixed, and compared three short-read-only MH63 whole-genome runs (Supplemental Figures S4 and S2). For Figure S4, the frozen final calls of RUN1_V4, RUN2_V4.1 and RUN3_V4.1 were independently exported under the same SJ policy; the manually curated annotation was used only after export. All three runs used the same 19,928 AI cards, system prompt, temperature 0 and non-reasoning setting. RUN1_V4 used the deepseek-chat alias (V4 Flash) and 60 workers; RUN2_V4.1 and RUN3_V4.1 used deepseek-flash (V4.1 Flash) and 20 workers. The two V4.1 runs are independent API calls under the same recorded settings; the V4/V4.1 comparison is one cross-version observation. Gene gain and loss were counted against reference genes exact in, or missed by, the current annotation (Figure 1D).

For quality assessment across 16 genomes (Figure 1E and Supplemental Figure S5), proteins were translated from each paired annotation and genome with gffread v0.12.6; the longest translated isoform per gene was retained. BUSCO v4.1.4 was run in offline protein mode with poales_odb10 for grasses and eudicots_odb10 for the other species. OMAmer v2.1.2 and OMArk v0.5.0 used the May 2026 Viridiplantae database and species taxonomy identifiers. Changes were calculated as SJ or SNM minus the corresponding original annotation, in percentage points, and averaged without weighting the 16 genomes. These measures are protein-set quality proxies rather than gene-structure accuracy against independent reference annotations. Separately, TAIR12 original annotation served as the fixed reference for exact-structure retention in *Arabidopsis* (Supplemental Figure S1). The three MH63 IGV examples in Supplemental Figure S7 used the original, GeneArbiter and manually curated reference tracks with root, stamen and young-panicle RNA-seq views; display regions and gene-ID mappings are listed in Source_Data/Figure_S7_source_data.tsv.

#### Structural evidence checks for Figure 1G and 1H

Exact splice-junction detection was assessed across 25 annotation-independent, genome-only HISAT2 short-read runs by querying the same merged junction set for each model-defined coordinate. For the merge case in Figure 1H, the cross-gene junction Chr01:20462823-20473047:+ was independently checked in regional BAMs: 18,810 primary short-read alignments supported the exact junction, including 16,884 with MAPQ 60 and NH=1; 11,552 also had at least 20-bp anchors on both sides. Five flag-leaf long-read runs yielded 197 reads spanning this intron, 43 with all seven exact junctions and 28 covering the complete coding span. RepeatMasker annotations covered 6,506/10,225 bp of the intron, while 100-bp splice-boundary flanks had only locus-specific full-length matches at identity ≥90% and query coverage ≥80%, and adjacent 50-bp flanks did so at 100% identity and coverage. Protein evidence cards identified distinct near-full-length homologues for the split components in Figure 1G. For Figure 1H, BLASTP against Nipponbare protein Os01t0527400-01 gave 87.2% and 100% query coverage for the 625-aa reference and 644-aa merged proteins, respectively. The 77-aa reference protein partially matched OB01G27860.1 at 65.52% identity and 75.32% query coverage. ANNEVO and Helixer reproduced the merged CDS chain in Figure 1H; Tiberius predicted split models. These checks support the merged major coding transcript without excluding additional shorter transcripts.

### Supplemental figure legends

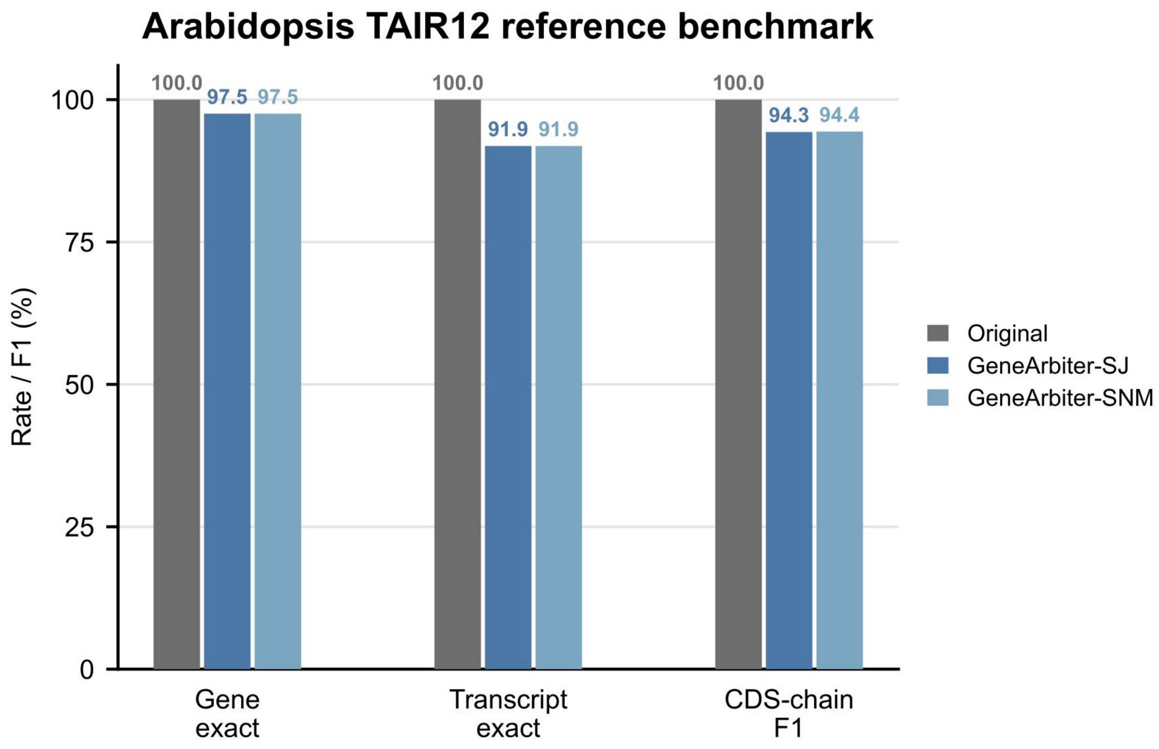


**Supplemental Figure S1. Retention of the original TAIR12 coding annotation by GeneArbiter-SJ and GeneArbiter-SNM.** The original TAIR12 annotation was used as the fixed reference for all three comparisons. Gene exact is the proportion of its 26,867 protein-coding genes for which at least one reference transcript has an exactly matched full CDS chain in the evaluated annotation. Transcript exact is the proportion of its 32,589 coding transcripts with an exactly matched full CDS chain; matches require the same chromosome and strand. CDS-chain F1 combines precision and recall for exact full-chain matches. Both GeneArbiter outputs matched 26,205/26,867 genes (97.54%) and 29,936/32,589 transcripts (91.86%); CDS-chain F1 was 94.32% for SJ and 94.38% for SNM. Original scores 100% for all three measures because it is compared with itself. The comparison measures retention relative to the original TAIR12 annotation; the remaining 662/26,867 genes lack an exactly retained full CDS chain, setting a conservative upper bound of 2.46% on potential unintended changes.

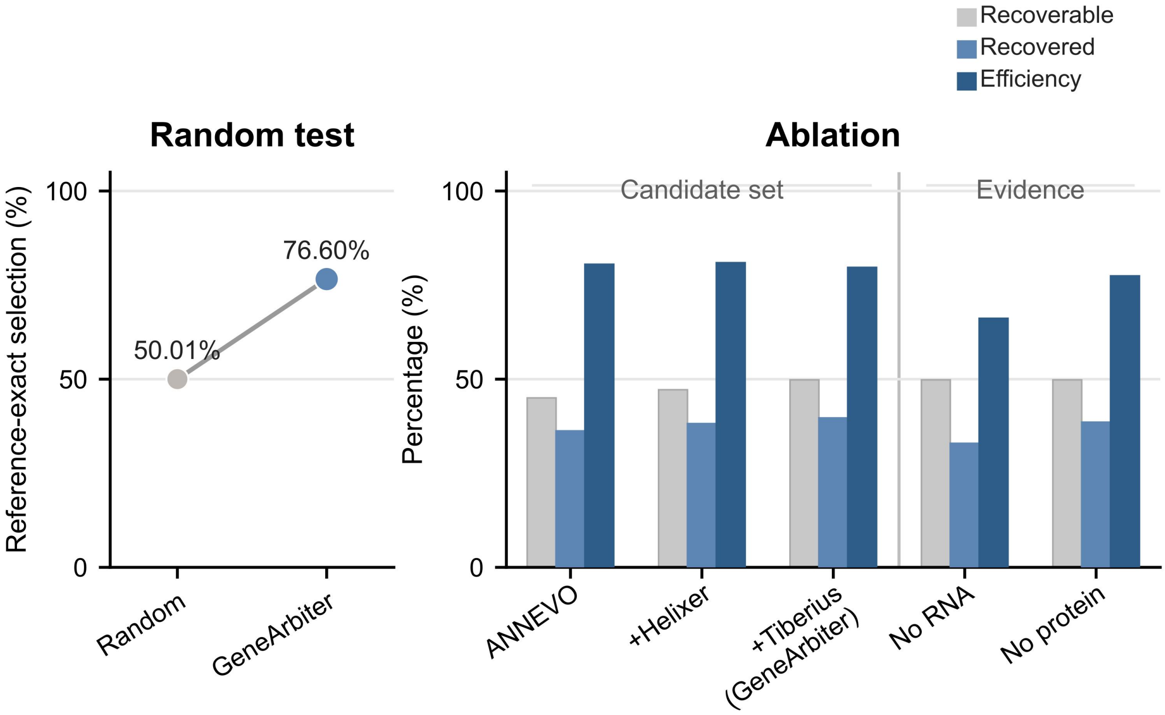


**Supplemental Figure S2. Reference-exact candidate selection and incremental candidate-set and evidence ablations in MH63.** A reference-exact candidate has a complete CDS-chain match to the manually curated MH63 annotation. Left, among 9,874 informative candidate cards with final calls and both reference-exact and non-exact candidate sets, GeneArbiter selected a reference-exact set in 76.60% of cards, compared with a 50.01% random expectation. For each card, the random expectation is the number of reference-exact candidate sets divided by all candidate sets; the plotted value is the mean across cards, not the outcome of a simulated random run. Right, a separate, reference-blind 500-gene sample enriched for alternative candidate structures was used for incremental ablation. The first three conditions added ANNEVO, Helixer and Tiberius in sequence to the current annotation's candidate pool. The last two retained the three-tool pool but omitted RNA or protein evidence, respectively. Among loci at which the current annotation differed from the curated reference, let $N$ be the number evaluated, $R$ the number with a reference-exact candidate, and $A$ the number whose final selection recovered a reference-exact candidate. Recoverable, recovered and efficiency denote $R/N$, $A/N$ and $A/R$, respectively. For ANNEVO, +Helixer, +Tiberius, No RNA and No protein, the ($N$, $R$, $A$) counts were (251, 113, 91), (267, 126, 102), (267, 133, 106), (267, 133, 88) and (267, 133, 103), respectively. The left and right panels use different analysis sets; the three ablation measures have different denominators and are not interchangeable measures of genome-wide accuracy.

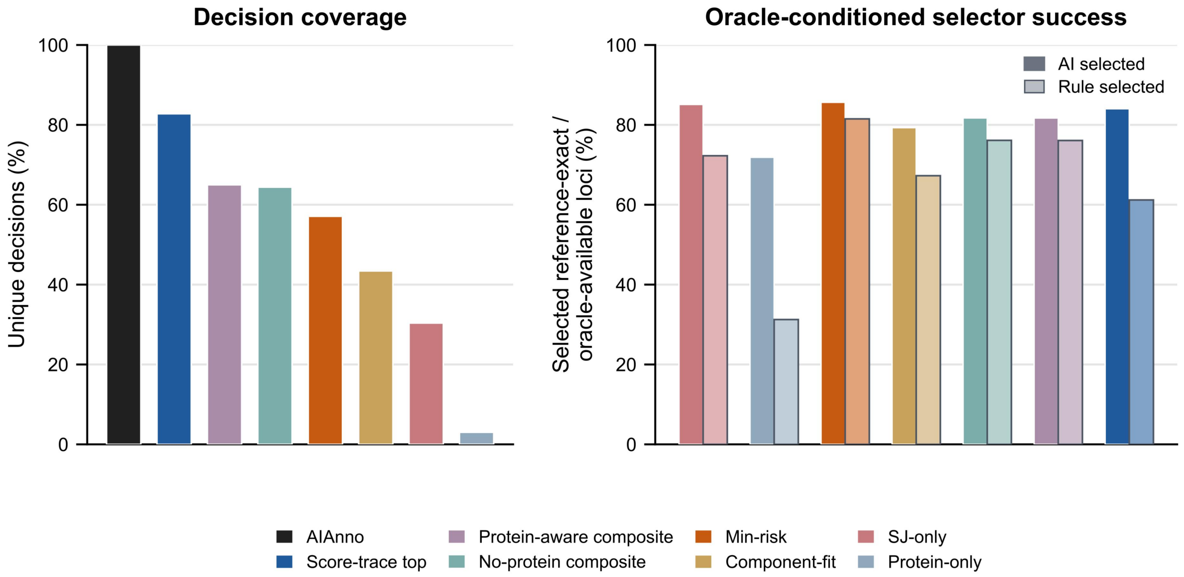


**Supplemental Figure S3. Constrained language-model arbitration compared with seven deterministic selectors in MH63.** The comparison used 19,928 API-routed loci with the same fixed candidate groups. Left, the proportion of loci receiving a unique selection; ties were counted as no-calls for deterministic selectors, whereas the constrained language model was required to return a listed candidate set. Right, reference-exact selection by the language model and each deterministic selector at loci where their choices differed and at least one reference-exact candidate was available. Each selector therefore has its own comparison set and denominator. The deterministic selectors used no curated-reference labels during selection. Source Data provide the counts and denominators for both panels.

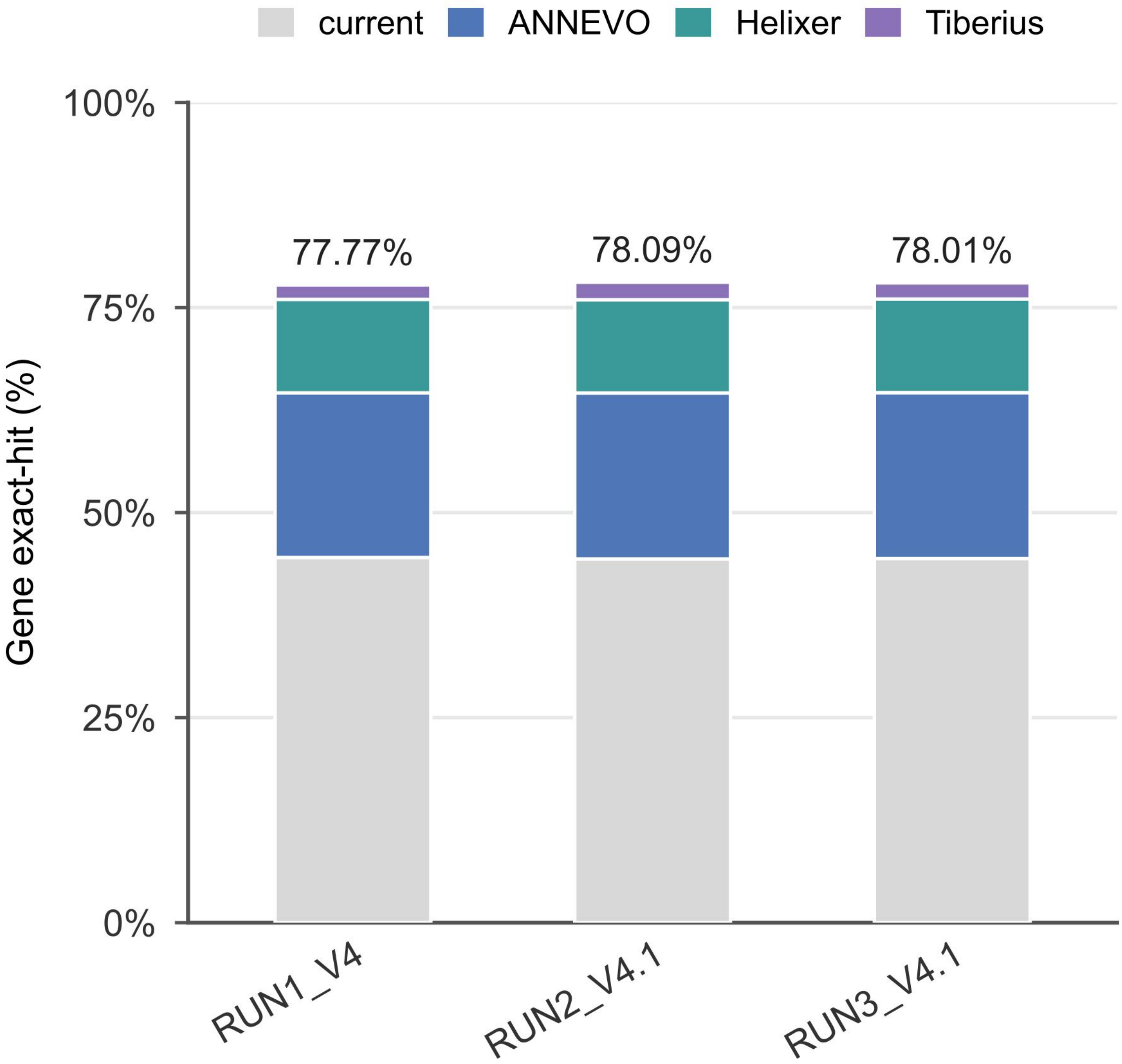


**Supplemental Figure S4. Gene-level exact-hit rates and selected-source composition across three whole-genome MH63 runs.** Each stacked bar gives the gene-level exact-hit rate after the same SJ release workflow: RUN1_V4, 31,216/40,140 (77.77%); RUN2_V4.1, 31,346/40,140 (78.09%); and RUN3_V4.1, 31,315/40,140 (78.01%). A reference coding gene is counted when at least one predicted transcript has an exactly matching complete CDS interval chain on the same chromosome and strand. Colours show the proportions of 43,530 frozen final calls selecting the original annotation, ANNEVO, Helixer or Tiberius, scaled to each bar's height; a coloured segment does not represent that source's contribution to exact hits. RUN2_V4.1 and RUN3_V4.1 used the same 19,928 AI cards, system prompt, temperature 0 and non-reasoning setting and selected the same candidate set in 19,500 shared valid API cards (97.85%). The V4/V4.1 comparison is a single cross-version observation. Source Data provide the plotted counts, denominators and segment heights.

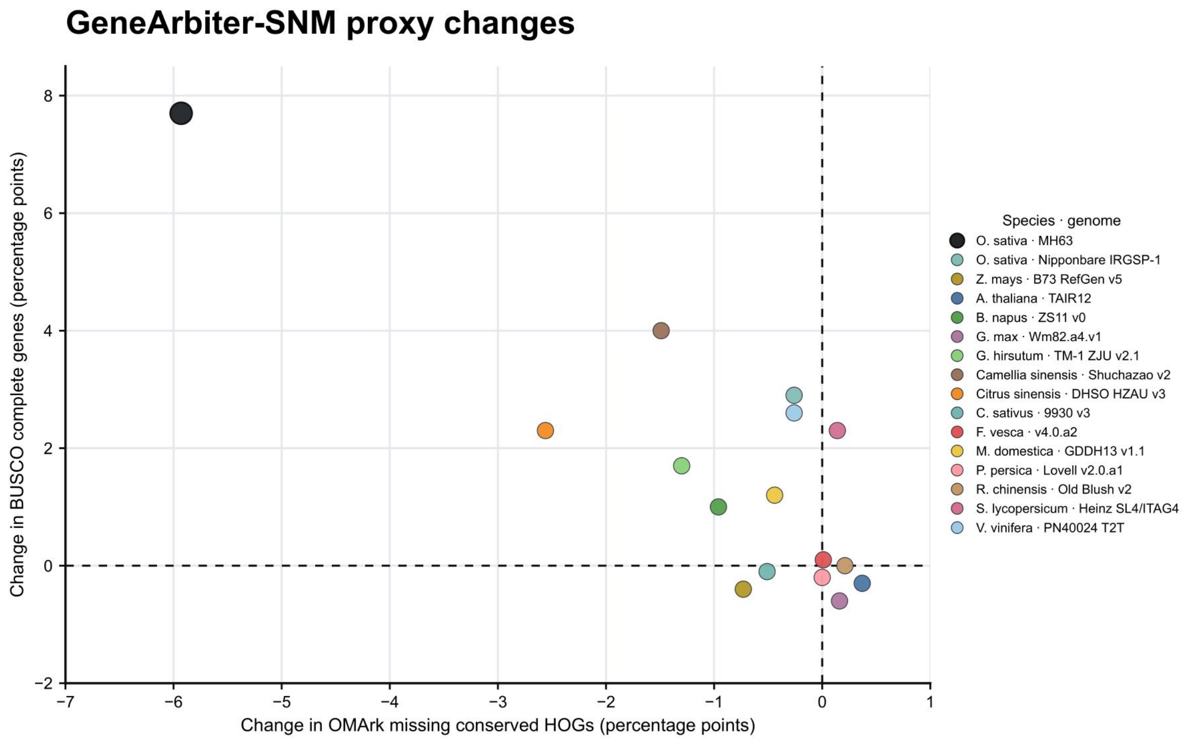


**Supplemental Figure S5. Paired changes in BUSCO and OMArk quality metrics under the conservative GeneArbiter-SNM release policy across 16 plant genomes.** Each coloured point represents one genome, with its original annotation as the paired baseline. For each annotation, the longest translated protein per gene was used for evaluation. SNM is the conservative release subset in which newly accepted coding structures require exact CDS-chain support from at least two independent non-current predictor sources. The vertical axis shows the change in BUSCO completeness, and the horizontal axis the change in the proportion of missing conserved hierarchical orthologous groups (HOGs) reported by OMArk; both are SNM minus original in percentage points. Dashed lines mark zero change. A positive vertical value or a negative horizontal value indicates improvement in the corresponding metric. Across the 16 genomes, the unweighted mean changes were +1.513 and −0.847 percentage points, respectively. BUSCO completeness increased in 10 genomes, was unchanged in one and decreased in five; OMArk missing HOGs decreased in 10, were unchanged in one and increased in five. BUSCO and OMArk reflect changes in quality metrics relative to the original annotations.

A

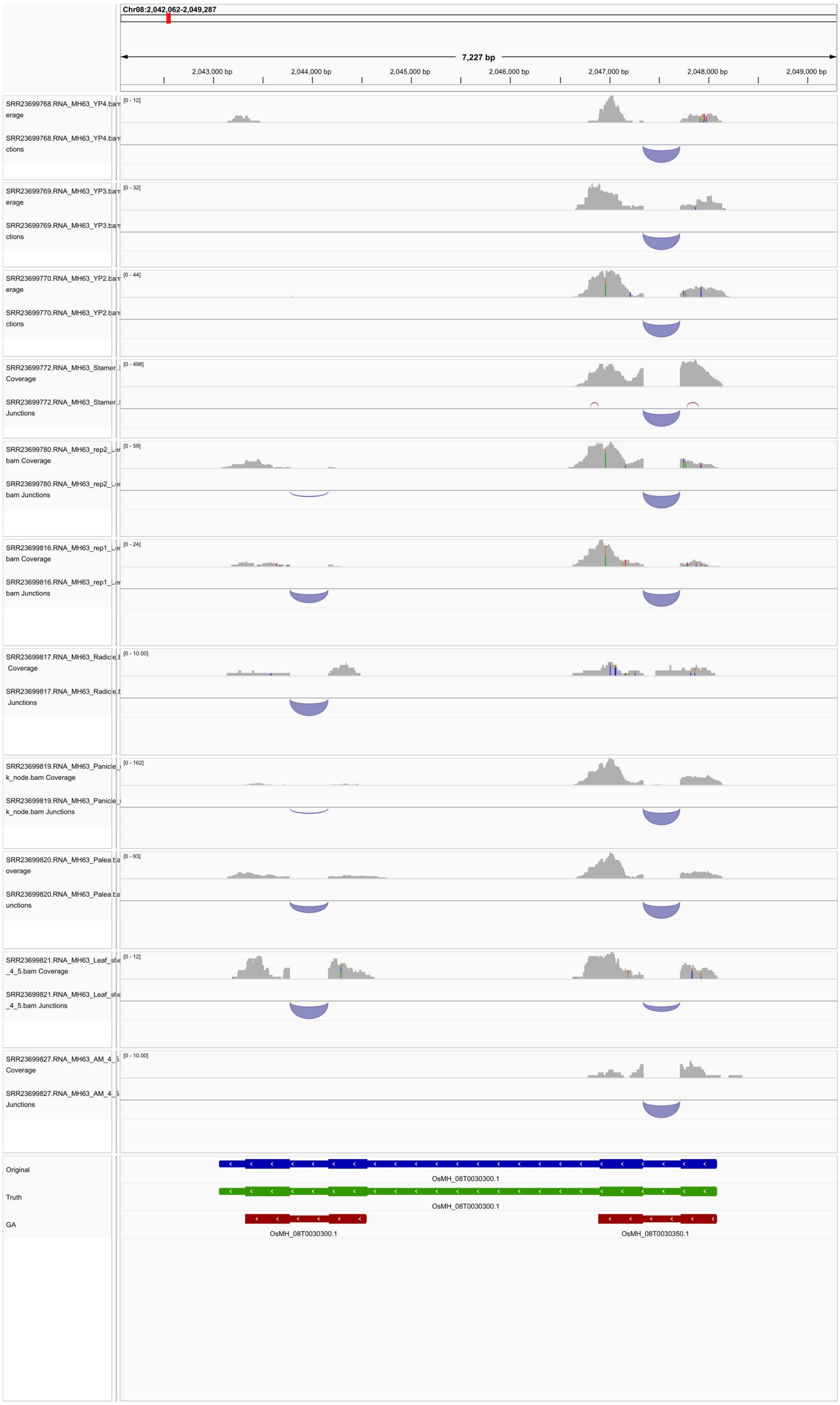

Chr08:2,042,062-2,049,287
7,227 bp
2,043,000 bp
2,044,000 bp
2,045,000 bp
2,046,000 bp
2,047,000 bp
2,048,000 bp
2,049,000 bp
SRR23699768.RNA_MH63_YP4.bam Coverage
SRR23699768.RNA_MH63_YP4.bam Junctions
[0 - 12]
SRR23699769.RNA_MH63_YP3.bam Coverage
SRR23699769.RNA_MH63_YP3.bam Junctions
[0 - 32]
SRR23699770.RNA_MH63_YP2.bam Coverage
SRR23699770.RNA_MH63_YP2.bam Junctions
[0 - 44]
SRR23699772.RNA_MH63_Stamen Coverage
SRR23699772.RNA_MH63_Stamen Junctions
[0 - 498]
SRR23699780.RNA_MH63_rep2_ bam Coverage
SRR23699780.RNA_MH63_rep2_ bam Junctions
[0 - 59]
SRR23699816.RNA_MH63_rep1_ bam Coverage
SRR23699816.RNA_MH63_rep1_ bam Junctions
[0 - 24]
SRR23699817.RNA_MH63_Radicle Coverage
SRR23699817.RNA_MH63_Radicle Junctions
[0 - 10.00]
SRR23699819.RNA_MH63_Panicle_ k_node.bam Coverage
SRR23699819.RNA_MH63_Panicle_ k_node.bam Junctions
[0 - 162]
SRR23699820.RNA_MH63_Palea Coverage
SRR23699820.RNA_MH63_Palea Junctions
[0 - 93]
SRR23699821.RNA_MH63_Leaf_ _4_5.bam Coverage
SRR23699821.RNA_MH63_Leaf_ _4_5.bam Junctions
[0 - 12]
SRR23699827.RNA_MH63_AM_4_ Coverage
SRR23699827.RNA_MH63_AM_4_ Junctions
[0 - 10.00]
Original
OsMH_08T0030300.1
Truth
OsMH_08T0030300.1
GA
OsMH_08T0030300.1
OsMH_08T0030350.1

**B**

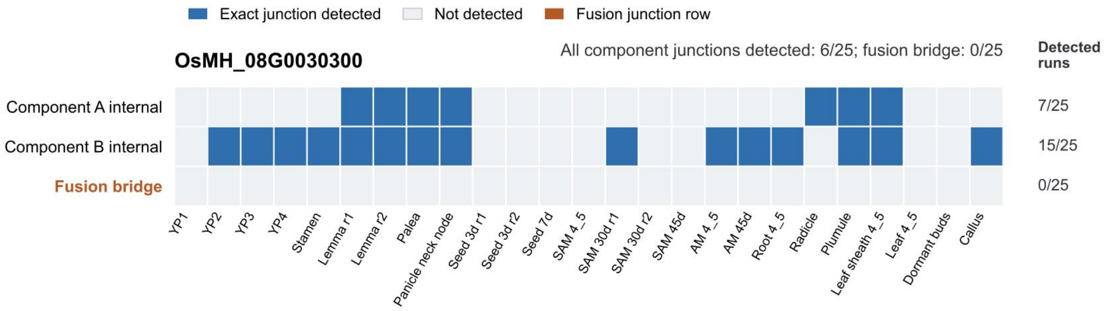
Exact junction detected
Not detected
Fusion junction row
OsMH_08G0030300
All component junctions detected: 6/25; fusion bridge: 0/25
Detected runs
Component A internal
Component B internal
Fusion bridge
7/25
15/25
0/25
YP1
YP2
YP3
YP4
Stamen
Lemma r1
Lemma r2
Palea
Panicle neck node
Seed 3d r1
Seed 3d r2
Seed 7d
SAM 4_5
SAM 30d r1
SAM 30d r2
SAM 45d
AM 4_5
AM 45d
Root 4_5
Radicle
Plumule
Leaf sheath 4_5
Leaf 4_5
Dormant buds
Callus

**C**

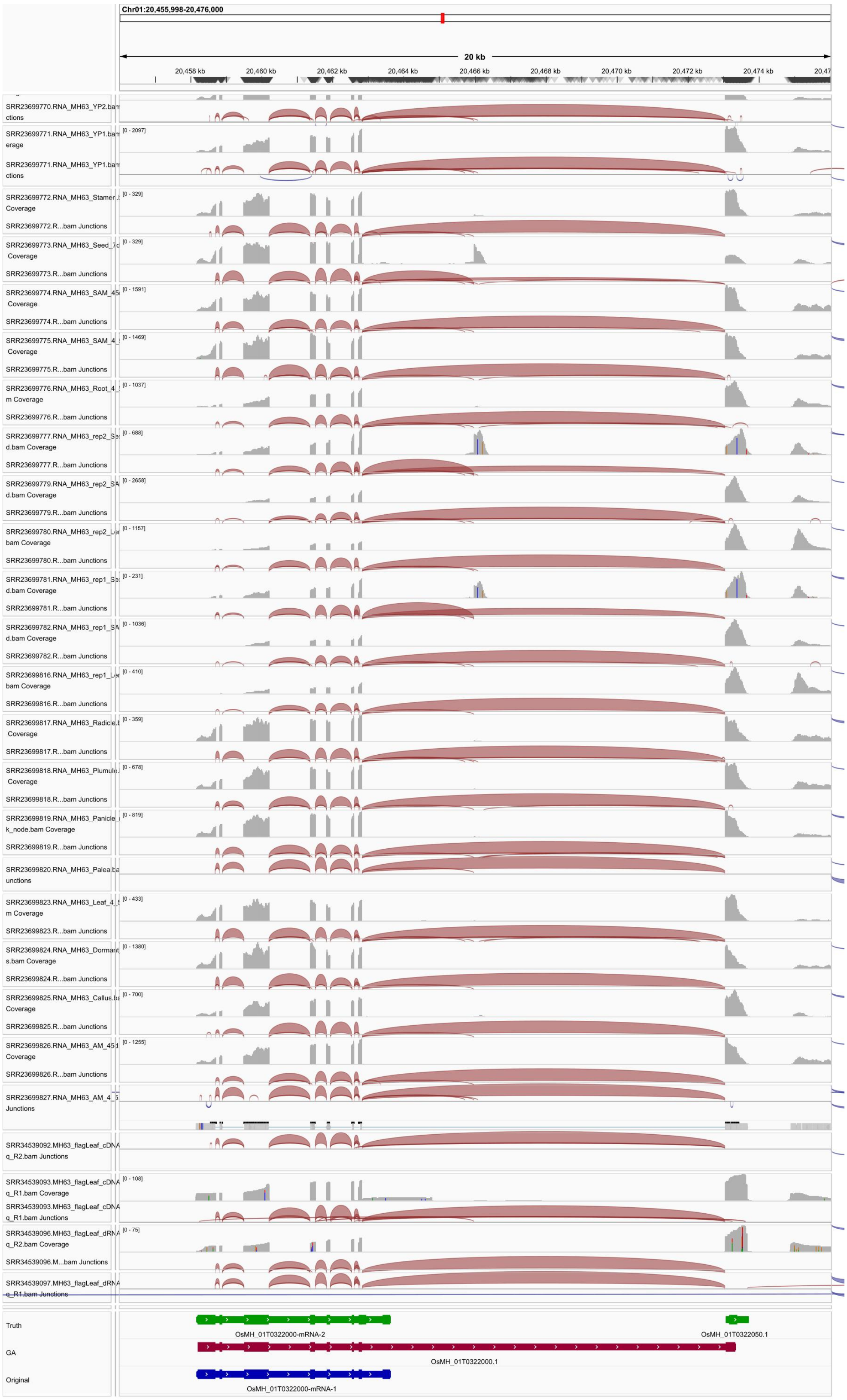

Chr01:20,455,998-20,476,000
20 kb
20,458 kb
20,460 kb
20,462 kb
20,464 kb
20,466 kb
20,468 kb
20,470 kb
20,472 kb
20,474 kb
SRR23699770.RNA_MH63_YP2.ba
ctions
SRR23699771.RNA_MH63_YP1.ba
erage
[0 - 2097]
SRR23699771.RNA_MH63_YP1.ba
ctions
SRR23699772.RNA_MH63_Stamen
Coverage
[0 - 329]
SRR23699772.R...bam Junctions
SRR23699773.RNA_MH63_Seed_7
Coverage
[0 - 329]
SRR23699773.R...bam Junctions
SRR23699774.RNA_MH63_SAM_45
Coverage
[0 - 1591]
SRR23699774.R...bam Junctions
SRR23699775.RNA_MH63_SAM_4_
Coverage
[0 - 1469]
SRR23699775.R...bam Junctions
SRR23699776.RNA_MH63_Root_4_
m Coverage
[0 - 1037]
SRR23699776.R...bam Junctions
SRR23699777.RNA_MH63_rep2_S
d.bam Coverage
[0 - 688]
SRR23699777.R...bam Junctions
SRR23699779.RNA_MH63_rep2_S
d.bam Coverage
[0 - 2658]
SRR23699779.R...bam Junctions
SRR23699780.RNA_MH63_rep2_L
bam Coverage
[0 - 1157]
SRR23699780.R...bam Junctions
SRR23699781.RNA_MH63_rep1_S
d.bam Coverage
[0 - 231]
SRR23699781.R...bam Junctions
SRR23699782.RNA_MH63_rep1_S
d.bam Coverage
[0 - 1036]
SRR23699782.R...bam Junctions
SRR23699816.RNA_MH63_rep1_L
bam Coverage
[0 - 410]
SRR23699816.R...bam Junctions
SRR23699817.RNA_MH63_Radicle
Coverage
[0 - 359]
SRR23699817.R...bam Junctions
SRR23699818.RNA_MH63_Plumule
Coverage
[0 - 678]
SRR23699818.R...bam Junctions
SRR23699819.RNA_MH63_Panicle
k_node.bam Coverage
[0 - 819]
SRR23699819.R...bam Junctions
SRR23699820.RNA_MH63_Palea.b
unctions
SRR23699823.RNA_MH63_Leaf_4_
m Coverage
[0 - 433]
SRR23699823.R...bam Junctions
SRR23699824.RNA_MH63_Dormant
s.bam Coverage
[0 - 1380]
SRR23699824.R...bam Junctions
SRR23699825.RNA_MH63_Callus.b
Coverage
[0 - 700]
SRR23699825.R...bam Junctions
SRR23699826.RNA_MH63_AM_45
Coverage
[0 - 1255]
SRR23699826.R...bam Junctions
SRR23699827.RNA_MH63_AM_4_
Junctions
SRR34539092.MH63_flagLeaf_cDNA
q_R2.bam Junctions
SRR34539093.MH63_flagLeaf_cDNA
q_R1.bam Coverage
[0 - 108]
SRR34539093.MH63_flagLeaf_cDNA
q_R1.bam Junctions
SRR34539096.MH63_flagLeaf_dRNA
q_R2.bam Coverage
[0 - 75]
SRR34539096.M...bam Junctions
SRR34539097.MH63_flagLeaf_dRNA
q_R1.bam Junctions
Truth
OsMH_01T0322000-mRNA-2
OsMH_01T0322050.1
GA
OsMH_01T0322000.1
Original
OsMH_01T0322000-mRNA-1

**D**

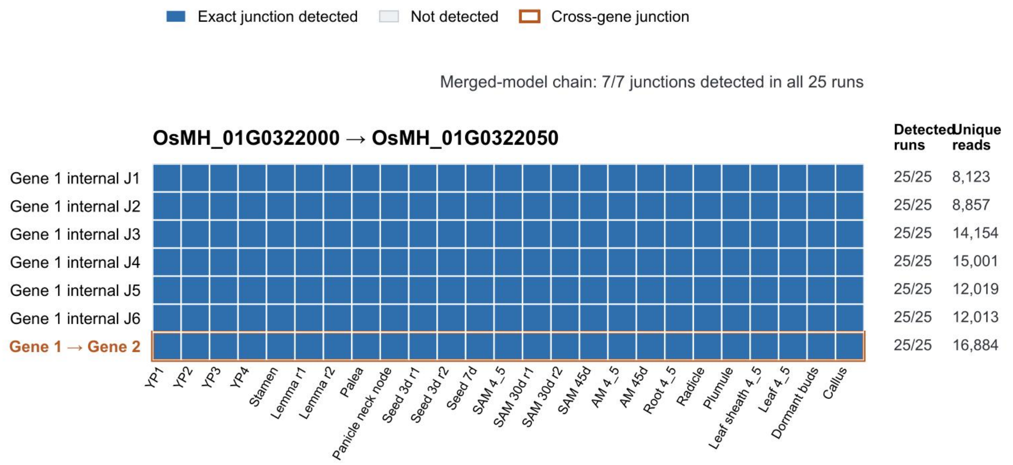


**Supplemental Figure S6. Structural evidence at two MH63 loci where GeneArbiter differs from the manually refined reference annotation. A, B,** At OsMH_08G0030300, the reference annotation represents one fused gene, whereas GeneArbiter selects two adjacent coding models. A shows selected RNA-seq coverage and splice-junction tracks together with the original, reference and GeneArbiter models. B shows exact-junction detection across all 25 short-read RNA-seq runs: the two component-internal junctions are detected in 7 and 15 runs, respectively, while the fusion junction is not detected in any run. The two split components match distinct near-full-length *Oryza glaberrima* proteins at 92.86% and 98.18% identity. **C, D,** At OsMH_01G0322000/OsMH_01G0322050, the reference annotation separates two genes, whereas GeneArbiter selects a model connected across a 10.2-kb intron. C shows the annotation models and selected short- and long-read RNA-seq tracks. D shows that all seven junctions of the merged model, including the cross-gene junction, are detected in all 25 short-read runs. Long-read analysis found 197 reads spanning the cross-gene intron, including 43 containing all seven model junctions. BLASTP against the same 644-aa Nipponbare protein (Os01t0527400-01) gave 87.2% query coverage for the 625-aa reference protein and 100% for the 644-aa merged protein. In B and D, blue denotes exact-junction detection, pale grey denotes no detection, and orange marks the junction that distinguishes the competing structures. The IGV panels show selected tracks, whereas the heatmaps summarize the tested junctions across all 25 short-read runs.

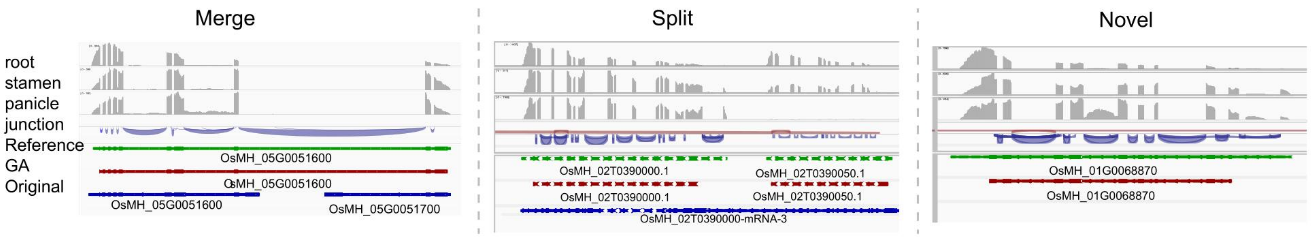


**Supplemental Figure S7. Representative MH63 loci showing three classes of GeneArbiter structural revision.** Left, merger of the original OsMH_05G0051600 and OsMH_05G0051700 models into OsMH_05G0051600. Middle, split of the original OsMH_02G0390000 model into OsMH_02G0390000 and OsMH_02G0390050. Right, addition of the current-absent OsMH_01G0068870 gene. IGV views show the original annotation, GeneArbiter model, manually curated reference and RNA-seq tracks from root, stamen and young panicle. Display regions and identifier mappings are provided in the Figure S7 Source Data file.

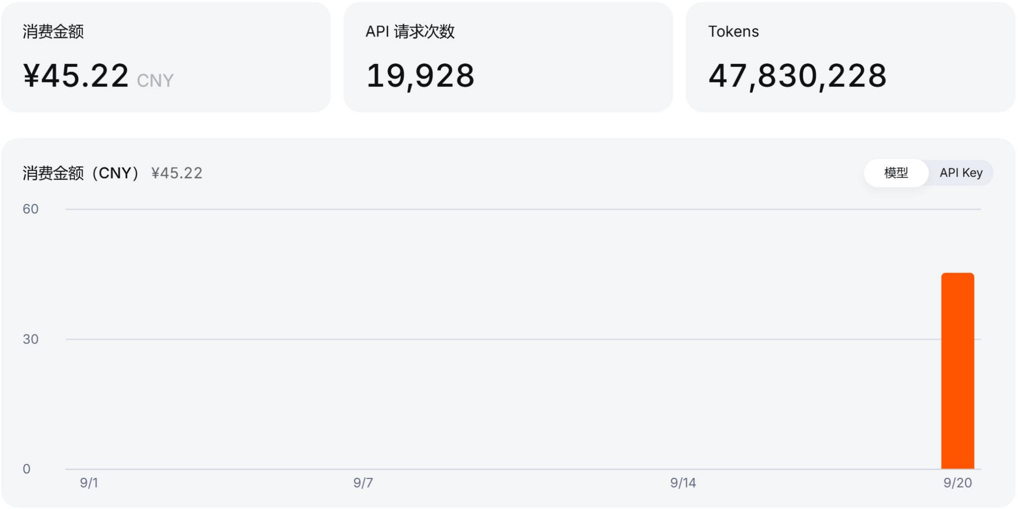


**Supplemental Figure S8. Observed API usage and charge for one MH63 V4.1 whole-genome run.** The provider dashboard screenshot shows CNY 45.22 in charges, 19,928 requests and 47,830,228 tokens. The request and token totals match the saved raw API responses for RUN2_V4.1, which comprise 44,536,716 prompt tokens (12,741,120 cache-hit and 31,795,596 cache-miss tokens) and 3,293,512 completion tokens. The charge is the displayed dashboard observation for this run; the screenshot does not report costs for the other 15 genomes. It represents direct API inference only, not the cost of candidate generation, evidence processing or local computation. Figure S8 Source Data report the matched counts; a separate Source Data TSV lists response-derived API token use for all 16 original whole-genome runs.

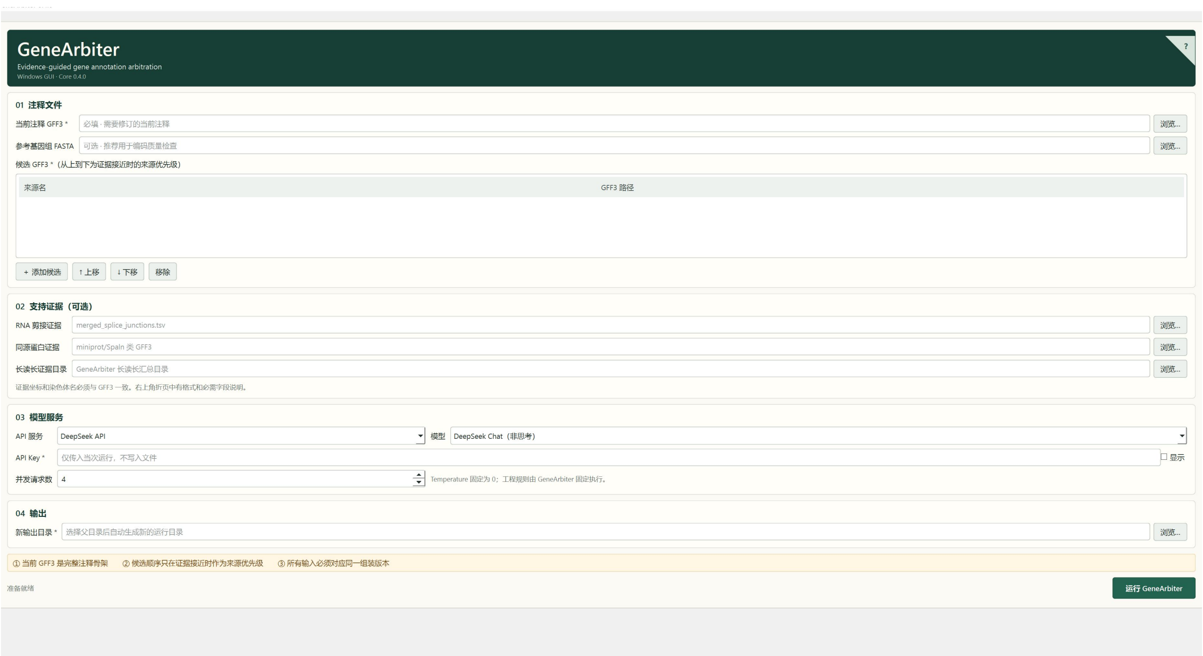


**Supplemental Figure S9A. GeneArbiter local graphical interface.** Screenshot of the Windows application configuration screen (Core 0.4.0, as displayed). The interface provides fields for the current GFF3 annotation, candidate GFF3 files, optional supporting evidence, model service settings and the output directory. This image shows the setup interface before an analysis is run.

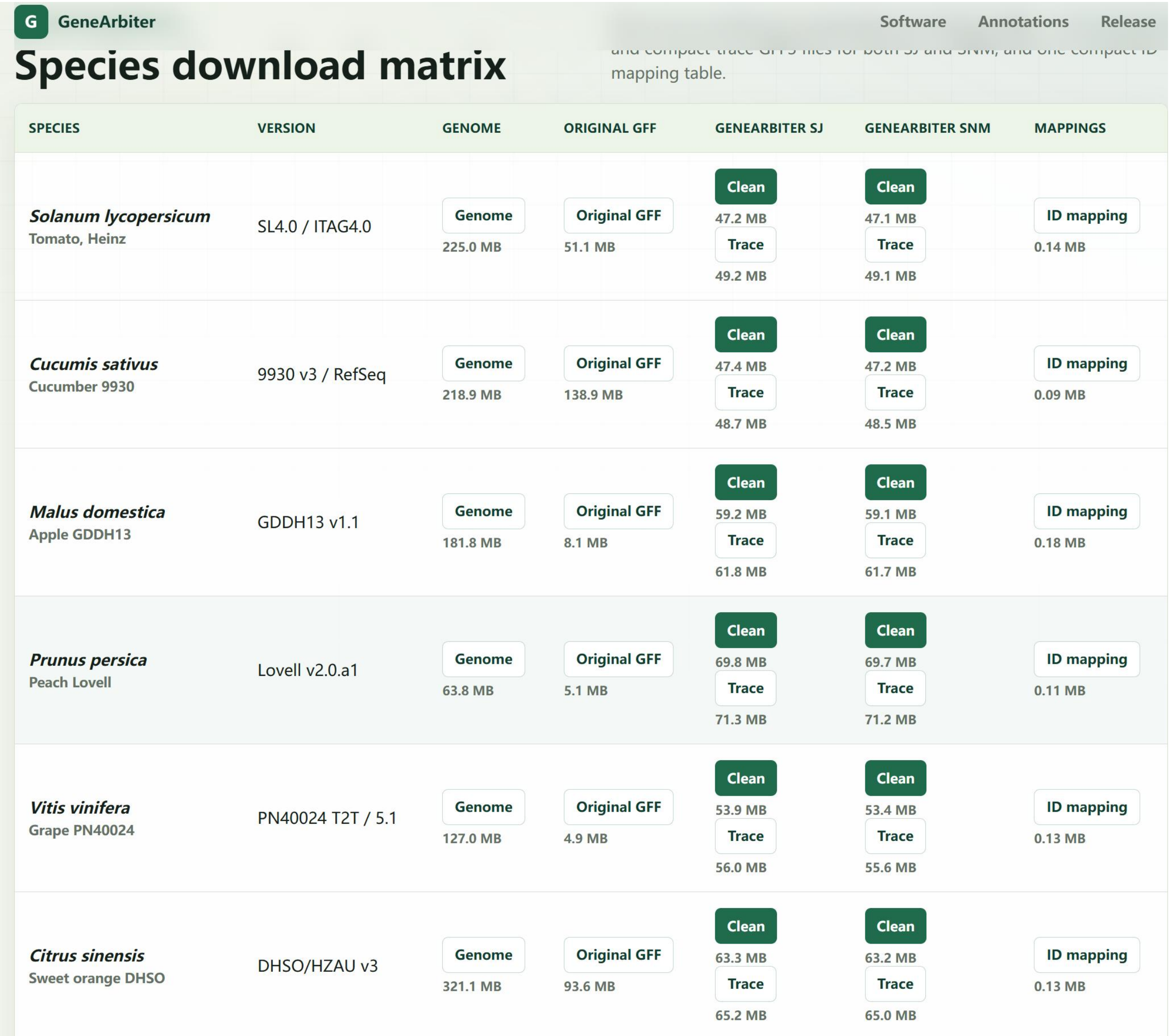


**Supplemental Figure S9B. GeneArbiter web portal.** Partial screenshot of the genome download matrix. The visible rows show links for genome assemblies, original GFF files, GeneArbiter-SJ and GeneArbiter-SNM annotation files, and gene identifier mappings. The screenshot illustrates the layout of the download interface; only part of the genome list is visible.

### Supplemental table legends

**Supplemental Table S1. Genome assemblies, baseline annotations and evidence resources for the 16-genome analysis.** Each row identifies one analysed genome, its baseline annotation and the RNA and homologous-protein evidence used. The two rice cultivars are separate genomes. The corresponding table is supplied in the accompanying spreadsheet.

**Supplemental Table S2. RNA-seq samples used for genome-wide evidence and the Figure 1G/H locus checks.** Rows identify the 113 selected samples, with tissue, developmental stage, replicate, library type, study role and public project accessions where available. The corresponding table is supplied in the accompanying spreadsheet.

**Supplemental Table S3. Software, resources and key settings.** Analysis steps, versions or identifiers, major parameters and outputs are listed in the accompanying spreadsheet.

#### Prompt appendix

The system message and per-card user template used for the MH63 short-read-only run are reproduced below. For each request, __CARD_JSON__ was replaced with the serialized compact card.

The following translation is provided for readers. The Chinese prompts below were used in the evaluated run; the English translation was not sent to the model.

##### System prompt

### English translation (for review only)

```
You are an adjudicator of gene-annotation evidence cards. Select an annotation set using only the AI annotation card supplied by the user.

General rules:
- Return a single JSON object only, without Markdown.
- Select only a set_id listed in card.sets; do not combine multiple sets.
- Do not generate coordinates, exons, CDSs, introns, transcripts, genes, or GFF records.
- By default, make a select_annotation_set decision. Use risk_flagged only when the card explicitly shows mutually exclusive candidate sets and the biological evidence cannot distinguish between them.
- Even for risk_flagged, provide a listed selected_set_id as the provisional best annotation set. risk_flagged is a label for manual review or export risk, not an abstention from selection.

Decision order:
1. Read decision_focus, set_comparison, and set_differences to identify the main conflict at this locus.
2. Read component_matrix, components, and each set's component_decision_summary to determine whether the differences concern a split or merger, a missing component, an extra component, an isoform or topology difference, or a coding risk.
3. For each set, jointly compare coding QC, phase consistency, splice and junction evidence, short-read support, long-read support, homologous-protein support, component support counts, and explicit risk tags.
4. Prefer a set with strong coding completeness, transcript and splice support, long-read support, and protein support that covers strongly supported components without an unsupported extra component.
5. Coding disruption, frameshifts, premature stops, phase inconsistency, unsupported or conflicting junctions, unsupported extra components, missing strongly supported components, and unsupported split or merger risks are important negative evidence.
6. candidate_priority does not prohibit a selection. However, low_priority_* means that an extra or missing component or a split or merger topology lacks direct support. Prefer such a set only when the other sets have weaker evidence or the structure has direct splice, long-read, or protein support.
7. Missing RNA, protein, or long-read evidence does not prove absence. It may reflect lack of expression, incomplete coverage, or an incomplete library and cannot by itself rule out a candidate.
8. Agreement among multiple candidate-generation sources indicates agreement between predictions, not biological correctness.
9. If the evidence does not distinguish the sets but the rules do not allow risk_flagged, select a listed set with low confidence and requires_validation=true.

Restrictions on risk_flagged:
- Use risk_flagged only when set_comparison.recommended_default is risk_if_unresolved_conflict and discriminator_status indicates conflicting or insufficient evidence.
- If recommended_default is select_best_listed_set, select a listed set_id.
- For risk_flagged, selected_set_id must still be a listed set_id; do not leave it empty.
```

### Chinese original (used in evaluation)

```
你是基因注释 evidence card 裁决器。你只根据用户提供的 AI annotation card 做 annotation-set 选择。

总规则：
- 只返回一个 JSON object，不要 markdown。
- 只能选择 card.sets 中已经列出的 set_id，不能混合多个 set。
- 不能生成坐标、exon、CDS、intron、transcript、gene 或 GFF 记录。
- 默认必须做 select_annotation_set；只有 card 明确显示候选 set 互相排斥，且生物学证据无法区分时，才允许 risk_flagged。
```

- 即使输出 risk_flagged，也必须同时给出一个 listed selected_set_id，表示 provisional best annotation set；risk_flagged 只是人工复核/导出风险标签，不是放弃裁决。

判断顺序：
1. 先读 decision_focus、set_comparison 和 set_differences，确定本 locus 的主要冲突。
2. 再读 component_matrix、components 和每个 set 的 component_decision_summary，判断差异来自 split/merge、component absence、extra component、isoform/topology 差异还是 coding risk。
3. 对每个 set，综合比较 coding QC、phase consistency、splice/junction evidence、short-read support、long-read support、homolog protein support、component support counts 和 explicit risk tags。
4. 优先选择同时具备较好 coding 完整性、转录本/剪接支持、long-read 支持、protein 支持，并覆盖强证据 component 且没有 unsupported extra component 的 set。
5. coding disruption、frameshift、premature stop、phase inconsistency、unsupported junction、conflicting junction、unsupported extra component、missing strong component、unsupported split/merge risk 是重要负面证据。
6. candidate_priority 不是禁止选择；但 low_priority_* 表示该 set 的额外 component、缺失 component 或 split/merge topology 缺少直接支持，只有其它 set 证据更差或该结构有直接 splice/long-read/protein 支持时才优先选择。
7. RNA、protein 或 long-read 缺失不是 absence 的证明，可能只是未表达、数据不覆盖或库不完整，不能单独作为否定证据。
8. 多个候选生成来源一致只代表候选一致性，不等于生物学正确性。
9. 如果没有足够证据区分，但规则不允许 risk_flagged，也必须选择一个 listed set，并使用 low confidence 和 requires_validation=true。

risk_flagged 使用限制：
- 只有 set_comparison.recommended_default 为 risk_if_unresolved_conflict，且 discriminator_status 显示证据冲突或无法区分时，才允许 risk_flagged。
- 如果 recommended_default 是 select_best_listed_set，必须选择一个 listed set_id。
- risk_flagged 时 selected_set_id 也必须是 listed set_id；不要留空。

#### User prompt template

### English translation (for review only)

```
Return a constrained decision for this AI annotation card.

Return a JSON object in the following format:
{
  "card_id": "copy card_id exactly",
  "decision": "select_annotation_set | risk_flagged",
  "selected_set_id": "must be a listed set_id; provide the provisional best set even for risk_flagged",
  "confidence": "high | medium | low",
  "reason_tags": ["short English tags"],
  "risk_tags": ["short English tags"],
  "evidence_summary": "one sentence explaining the evidence; cite only information in the card and do not introduce new coordinates",
  "requires_validation": true
}

AI annotation card:
__CARD_JSON__
```

### Chinese original (used in evaluation)

```
请为这个 AI annotation card 返回受约束决策。

必须返回 JSON object，格式如下：
{
  "card_id": "必须逐字复制 card_id",
  "decision": "select_annotation_set | risk_flagged",
  "selected_set_id": "必须是 listed set_id；risk_flagged 时也必须给 provisional best set",
  "confidence": "high | medium | low",
  "reason_tags": ["英文短标签"],
  "risk_tags": ["英文短标签"],
```

```
  "evidence_summary": "一句话说明依据，只能引用 card 内信息，不要给出新坐标",
  "requires_validation": true
}

AI annotation card:
__CARD_JSON__
```
